# Estimating the scale of biomedical data generation using text mining

**DOI:** 10.1101/182857

**Authors:** Gabriel Rosenfeld, Dawei Lin

## Abstract

While the impact of biomedical research has traditionally been measured using bibliographic metrics such as citation or journal impact factor, the data itself is an output which can be directly measured to provide additional context about a publication’s impact. Data are a resource that can be repurposed and reused providing dividends on the original investment used to support the primary work. Moreover, it is the cornerstone upon which a tested hypothesis is rejected or accepted and specific scientific conclusions are reached. Understanding how and where it is being produced enhances the transparency and reproducibility of the biomedical research enterprise. Most biomedical data are not directly deposited in data repositories and are instead found in the publication within figures or attachments making it hard to measure. We attempted to address this challenge by using recent advances in word embedding to identify the technical and methodological features of terms used in the free text of articles’ methods sections. We created term usage signatures for five types of biomedical research data, which were used in univariate clustering to correctly identify a large fraction of positive control articles and a set of manually annotated articles where generation of data types could be validated. The approach was then used to estimate the fraction of PLOS articles generating each biomedical data type over time. Out of all PLOS articles analyzed (n = 129,918), ~7%, 19%, 12%, 18%, and 6% generated flow cytometry, immunoassay, genomic microarray, microscopy, and high-throughput sequencing data. The estimate portends a vast amount of biomedical data being produced: in 2016, if other publishers generated a similar amount of data then roughly 40,000 NIH-funded research articles would produce ~56,000 datasets consisting of the five data types we analyzed.

**One Sentence Summary:** Application of a word-embedding model trained on the methods sections of research articles allows for estimation of the production of diverse biomedical data types using text mining.

## Introduction

### Biomedical data as a critical research output

Biomedical research is a fundamental component of the United States’ research and development portfolio and has led to numerous benefits both economically and scientifically. It is imperative to continually measure the outputs of this publicly funded scientific endeavor so stakeholders (Congress, the public, biomedical investigators, etc.) understand the impact of the tax dollars being spent. Most of these analyses, so far, have focused on the impact of the scientific work from a bibliographic lens to address questions such as how many times a paper resulting from the research was cited or the impact factor of the journal in which the paper was published(1). Recent innovations have included additional metrics such as patents while simultaneously making portfolio analysis and data integration easier(2) as well as measures to normalize citation across various scientific disciplines(3). While these approaches provide a useful means of determining the impact of a research project, there are other outputs that are also measurable and indicative of impact. One example is the biomedical data itself produced during the study. Such data can be found within the articles (as figures, attached excel files, images, etc.) or can be deposited separately within a dedicated repository (e.g. GEO, SRA, ImmPort, etc.) that is referenced to within the article. There is an opportunity to leverage this information as an important output of the research project.

Measuring production of data from articles is beneficial for several reasons. Data is a resource which can be repurposed and reused for additional scientific studies further enhancing the return on investment of the original work (4). There is a vibrant ecosystem of open-source software developers and hobbyists interested in accessing scientific data that can help unleash this potential (5). One example is Ropensci, a community-driven project developing a collection of R packages that permit access to a variety of data repositories storing scientific data (6). These communities can facilitate development of user-driven secondary data products such as interactive visualizations, enhancements in machine learning and artificial intelligence, or entirely new, ancillary datasets in a virtuous cycle of discovery by these communities and their collaborators (5, 7, 8). As technologies like Galaxy or Shiny facilitate web-based access of data-informed discoveries and tools to a broader audience including bench scientists and the public, the data can have an even larger impact scientifically and socially (9, 10). As data are repurposed, investigators should be acknowledged if their dataset is consistently reused and deemed valuable by the research community. A burgeoning appreciation for data is showcased by Nature’s *Scientific Data Journal* and the *Biomedical Data Journal* which focus solely on the publication and dissemination of valuable datasets.

Just as important as the idea of data as a valuable commodity for reuse is the fact that the data itself is the cornerstone upon which transparency and reproducibility of the biomedical scientific endeavor is built. It forms the basis for whether a scientific hypothesis being tested is accepted or rejected, which is crucial for other investigators looking to understand the conclusions of the study. Beyond its importance to other researchers, information regarding the generation of data in an article can inform funding bodies, publishers, repositories, and other key stakeholders in the biomedical data ecosystem about the scale and diversity of biomedical data being produced (11). Such information forms the basis of data-informed management by helping to determine the cost of housing or sharing data and how the biomedical data ecosystem is evolving over time (12).

### How to measure the scale biomedical data?

Since most biomedical datasets are not deposited in a particular biomedical repository rather contained in the article itself(13), how to identify whether biomedical research articles are producing certain kinds of data is not a trivial issue. Like the biological processes and mechanisms being studied, the resulting data is diverse and the terminology describing the data reflects this diversity. While key words and MESH terms are a principled approach to develop a rough estimate of the production of various data types, they rely on manual annotation which may miss emerging or relevant terms resulting in a more conservative estimate (14). To address this challenge, we developed a text-mining strategy that identifies articles producing the following types of biomedical data: high-throughput sequencing, flow cytometry, immunoassay (e.g. ELISA, ELISPOT, multiplex assays, etc.), genomic microarray, and microscopy data. We selected these five data types to prototype the approach on a smaller scale while retaining a diversity of the types of biomedical data that the research community produces. Our approach builds upon advances in word embedding by leveraging the Global Vectors for Word Representation (GloVe) algorithm to identify a set of terms highly representative of each type of biomedical data we analyzed (15). We use this information to cluster articles via the total signal associated with these terms at the article level. One can customize the approach for additional data types to approximate the generation of biomedical data of interest using the free text of research articles.

These estimates can help put a quantity on the volume of biomedical data being produced to better capture this important research output from publicly funded studies. Our analysis using 129,918 PLOS articles published between 2003 and 2016 found that 59,543 articles generated one or more of the five biomedical data types we tested. In total, we assessed that 81, 407 data sets from one of the five data types were produced with a mean of ~ 1.4 data types per each data-producing article. This analysis supports a vast amount of data being produced within a limited scope. It is likely that a larger amount of biomedical data is being produced given the focus on only five data types from one publisher. For instance, a search of PubMed to identify NIH funded research articles published in 2016 that were not reviews resulted in 91, 685 articles. If a similar scale of data from our analysis were being produced within this set of articles, then approximately 40,000 would produce roughly 56,000 datasets consisting of one of the five biomedical data types we analyzed. A search query to identify PubMed research articles that were not reviews published in 2016 resulted in 1, 160, 334 articles. If the proportions of articles producing each data type were the same as we observed in PLOS subset, then roughly 81000, 220000, 140000, 210000, and 70000 of PubMed research articles in 2016 would produce flow cytometry, immunoassay, genomic microarray, microscopy, and high-throughput sequencing data respectively.

## Methodology

### Text Mining Analyses

#### Manually annotated articles

Articles were manually annotated as producing one of the five data types if the methods section or figures mentioned the use of certain technologies associated with a data type (e.g. ELISA, ELISPOT, or multiplex for immunoassay data; FACS or flow cytometry for flow cytometry data) or whether samples that could produce such data were used (see supplementary info for the complete list of manually annotated articles, their annotations, and origins).

#### Generating the GloVe model

Methods sections obtained from 381,651 open-access PMC articles were preprocessed for subsequent analysis by converting text to lower case, removing all non-alphanumeric characters, and merging any duplicated space into single space. A term-co-occurrence matrix was calculated from the pre-processed methods sections using the text2Vec package removing terms occurring less than ten times. A Global Vectors for Word Representation (GloVe) model was created using text2Vec with the following parameters: word_vectors_size = 300, vocabulary = preprocessed vocab from methods section, shuffle = F, x_max = 10, learning_rate = 0.15. The GloVe model was fit on the term-co-occurrence matrix generated from the methods sections with the following parameters: n_iterations = 20.

#### Generating a representative vector for each data type

The word embedding model generated by the GloVe algorithm was used to define a representative vector for five biomedical data types: high-throughput sequencing, flow cytometry, microscopy, immunoassay, and genomic microarray. To generate these representative vectors, a function was developed that takes an input of seed terms. The positive and negative seed terms were manually identified for each data type based upon domain knowledge of the terminology. The comprehensive procedure and mathematics behind the generation of the vector for each data type can be found in the supplementary information.

#### Generating univariate data for clustering

Using the text2vec package, a term frequency–inverse document frequency (TF-IDF) matrix was created from the abstracts, methods, or full texts of PLOS articles. The TF-IDF matrices were filtered on the closest terms to the representative vector for each data type from the GloVe model. The TF-IDF values of each filtered matrix were multiplied by the l2 cosine similarity measures of the respective terms to the representative vector of the corresponding data type. The total weights of all terms were summed to generate an article level univariate statistic called the “termFreq”. The full procedure and mathematics behind the generation of the “termFreq” can be found in the supplementary information.

#### Clustering analysis

For all clustering, the Ckmeans.1d.dp algorithm from the Ckmeans.1d.dp was used with the following parameters: k = 2.

##### Optimizing the recall using positive control

To test the effect of varying the number of closest terms used as well as the stochastic nature of the function when identifying the data type vector, a positive control set of articles indicative of the production of each biomedical data type was identified. The supplementary information contains detailed instructions on how positive controls were identified. 1000 random PLOS articles were repeatedly sampled using bootstrap (n = 5) and seeded with a sample of positive control articles for each data type. “termFreq” vectors were generated from the abstract, methods, or full text of PLOS articles using a variable number of ranked terms and resulting weights generated by altering the set.seed parameter in the data type generator function. These different “termFreq” vectors were clustered. The cluster with the highest mean “termFreq” statistic was compared to the fraction of positive control articles within that sample to estimate the recall on the positive controls.

##### Optimizing the approach using manually annotated articles

1000 random PLOS articles were repeatedly sampled using bootstrap (n = 5) and supplemented with the manually annotated articles (n = 177). The sampled articles were then clustered using the “termFreq” vectors generated from the abstract, methods, or full text when varying the number of closest terms or the set.seed parameter in the function generating the data type vector. The cluster with the highest mean “termFreq” was compared to the manual annotation to determine the precision and recall for each type of data.

## Results and Discussion

### Generating a GloVe model from methods sections

To develop an approach that could identify the data being generated by the biomedical research community, we capitalized on advances in natural language processing (NLP) involving word embedding. Word embedding is a feature learning technique in which words are mapped to vectors of real numbers (16). We hypothesized that the approach should capture the technical and methodological features of biomedical terminology used in research articles. We could use this information to identify different types of data being produced within a biomedical research article. We chose the GloVe model due to its speed and similarity in terms of NLP results to other word embedding approaches such as word2vec (15). The GloVe model was trained on the methods section as this is the section of the article enriched in the technical and methodological terminology.

1,334,297 XML files from the PubMed Central^®^ open-access subset were downloaded. The XML files contain a sec-type XML tag that allows for specific sections of the article to be processed and retrieved. We used this metadata to process the XML files and retrieve the abstract, methods, and full-text for articles containing the tags for these sections. 381,651 method sections were parsed and used to develop a GloVe model containing a vector representation of words that reflect the scientific community’s use of technical and methodological terms (Fig. 1). The final GloVe model contains a list of 267, 851 words with word embedding vectors of length 300 and is available to download and reuse (supplementary info). While longer vectors might capture additional features, previous work showed that use of vectors of length 300 were effective for many NLP tasks (17).

**Fig. 1.**
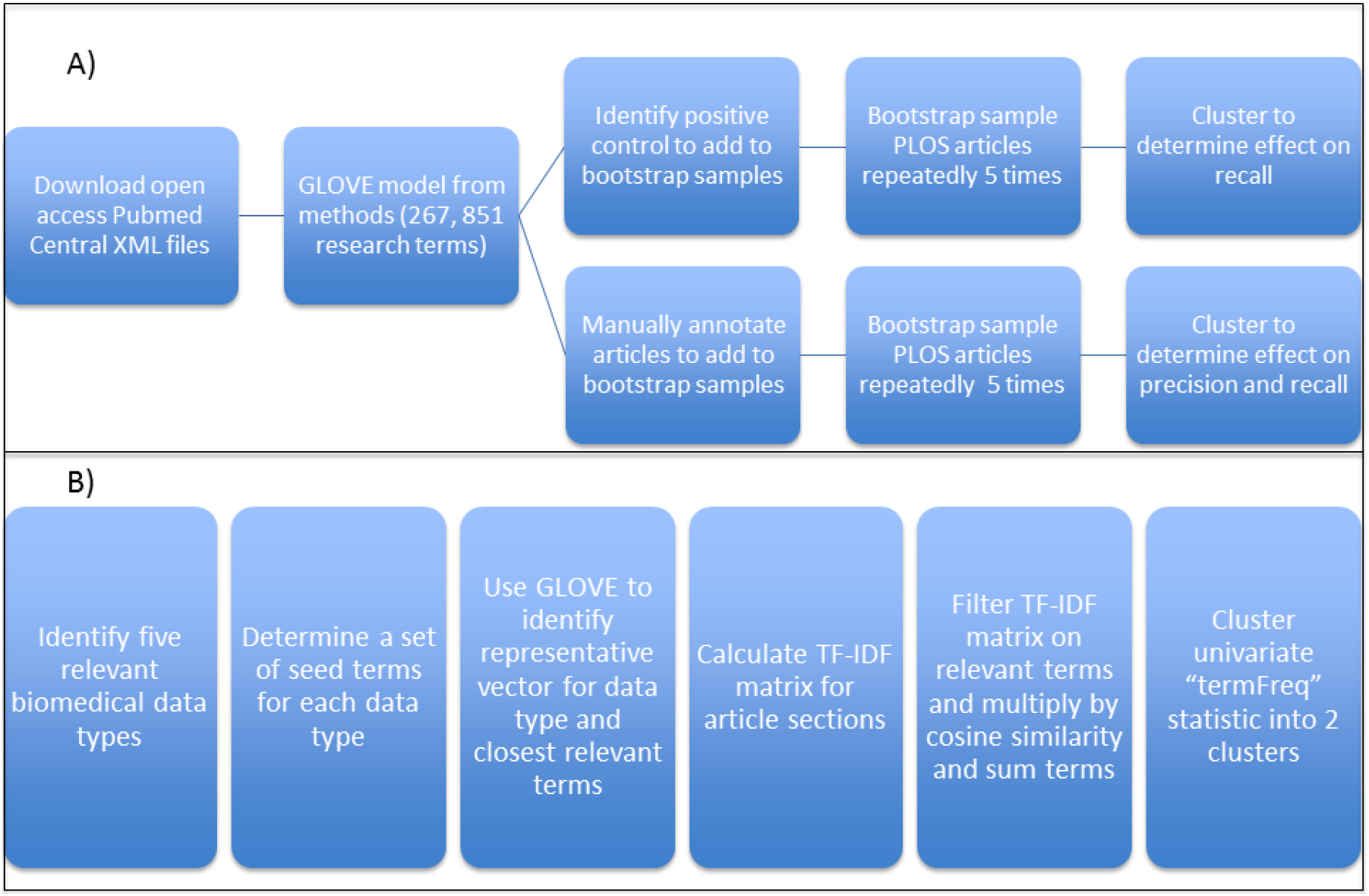
Overview of approach. An overview of the approach utilizing text mining to identify whether a research article produces five different types of biomedical data. A) The open access subset of PubMed Central articles was downloaded and a GLOVE model was trained on the parsed methods sections of these articles (n = 381,651). The GLOVE model identified the syntactic relationships of 267,851 terms and was used to generate five distinct word vectors identifying a set of terms indicative of a particular biomedical data type. Research articles were clustered into two subsets using the abstract, methods, or full text and the subset containing the higher fraction of the usage of particular biomedical terms was compared to positive control and manually annotated articles to determine the recall and precision of the approach. B) The clustering approach itself was tested on five data types (sequencing, microarray, immunoassay, flow cytometry, and microscopy) which represent a variety of biomedical data types of importance to immunology and are also present in the ImmPort database. Domain knowledge was used to identify a set of seed terms in order to use the GLOVE model to generate a word vector representing a particular data type. TF-IDF matrices were calculated using the different sections of the PLOS set of articles (abstract, methods, and full text) and subsequently filtered on the specific terms identified for a biomedical data type. The corresponding TF-IDF values were multiplied by l2 normalized cosine similarity of each term to the word vector for that data type. The resulting relevancy matrices were summed by each row to generate an article-level, univariate “termFreq” statistic, which was used to cluster the set of articles into two subsets using Ckmeans.1d.dp: one with low mean **“termFreq” and one with high mean “termFreq”. The high mean “termFreq” set** corresponds to the set of articles indicating generation of the particular data type.

### Identifying a list of words indicative of different types of biomedical data for clustering

We leveraged the GloVe model to develop a statistic at the article level that indicates generation of biomedical data. Five types of biomedical data were tested: high-throughput sequencing, genomic microarray, microscopy, immunoassay, and flow cytometry data. We selected this small number of terms to allow for fast prototyping of the approach while retaining a breadth of the diversity of data types that the biomedical research community produces. These data types are also curated in the Immunology Database and Analysis Portal, which NIAID DAIT funds so this analysis would provide data-driven insights directly of interest to the programmatic priorities of our Division (18).

We developed a function that takes a small set of input terms indicative of a data type and returns a full list of all terms associated with that data type sorted by their geometric distance to the data type using the word embedding properties from the GloVe model. These data type specific terms are used to generate a univariate statistic for clustering of articles into those with high usage of relevant terms and those with lower usage of less relevant terms. The function sums the vectors of the positive seed terms and subtracts the vectors of the negative terms to generate an input vector for a data type. Domain knowledge and empirical testing is required to identify the positive and negative terms to use. For instance, inputting terms for sequencing data resulted in terms for other related data types such as microarray and PCR data (so the terms, “microarray” and “pcr”, were subtracted from the input when generating the sequencing data type vector).

Iterative sampling of nearby terms to the input vectors (identified by l2 normalized cosine distance) is performed, the vectors for those terms are added to the input vector, and the nearest 10 terms of this new vector is compared to the 10 nearest terms of a summed vector of randomly sampled nearby terms. The iteration stops when the 10 nearest terms for these two vectors are equivalent or after a specified number of iterations. The goal is to generate a representative vector for each data type incorporating features of nearby terms by allowing the function to stochastically explore the feature space of these terms. The domain knowledge used to identify the terms can thereby be refined computationally to include potentially interesting features for a data type. A representative example of the top 100 terms identified for flow cytometry data is shown in Fig. 2. This visualization shows that the GloVe model is capturing the technical features of words from the methods section including the models of flow cytometers used, the manufacturers of the flow cytometers, the types of software used for the analyses, as well as the specific terminology used in flow cytometry experiments including the gating process. Similar properties are captured for the other data types (see supplementary information).

**Fig. 2.**
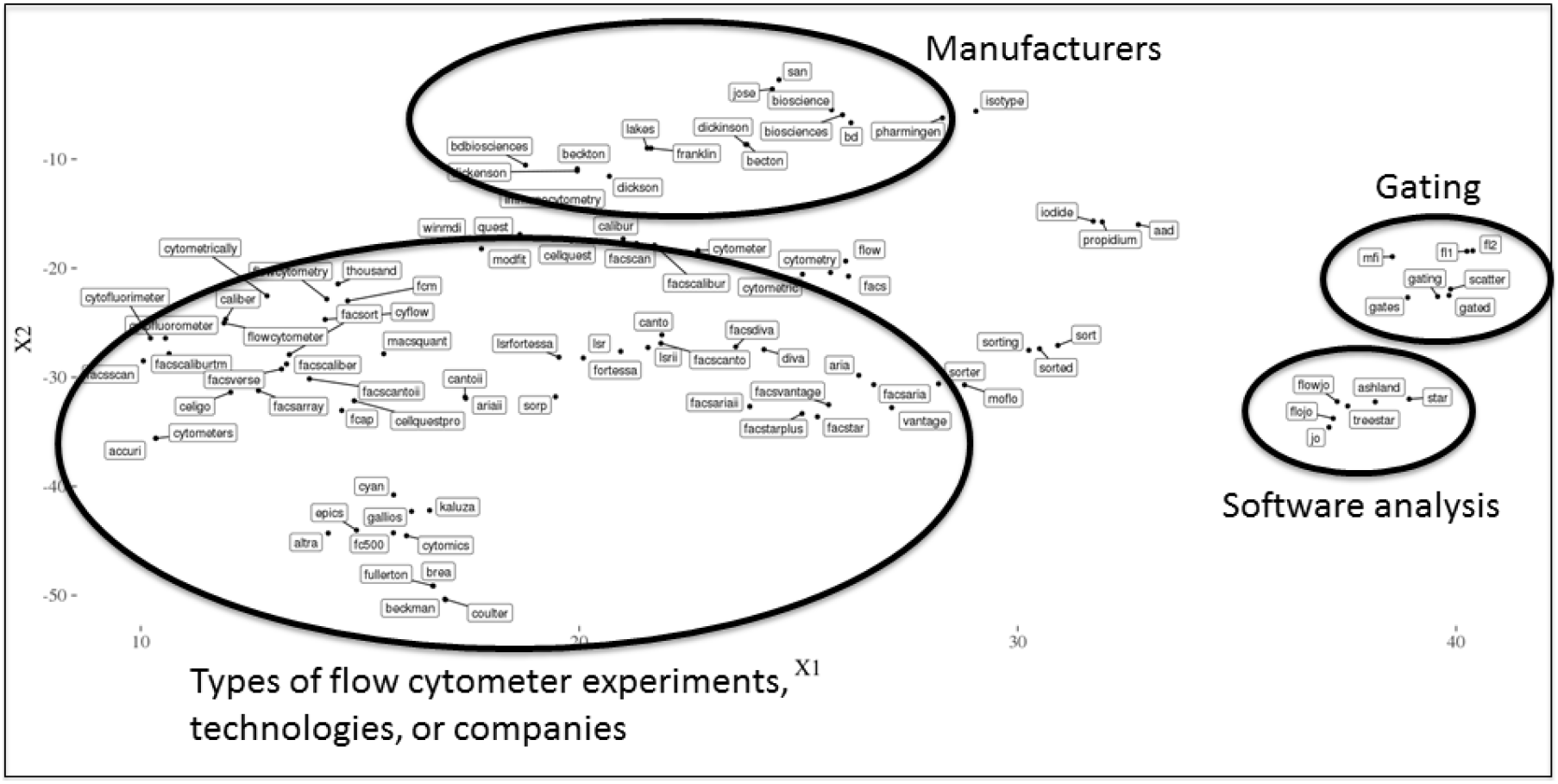
Flow cytometry data type top 100 terms. To give an illustration of the types of terms that are identified for a particular data type, the top 100 terms via l2 normalized cosine similarity to the flow cytometry data type word vector using optimized approach in Table 1 is shown. The word vectors for these terms were visualized using the T-SNE algorithm (using the Barnes-Hut approximation) to embed the high-dimensional vector relationships between the set of words into 2 dimensions (X1 and X2). The resulting visualization captures word similarities such that words often used in the same context within a methods section are nearer to each other. In this representative example for flow cytometry, one can observe the words clustering into smaller sub-clusters corresponding to specific themes like the specific flow cytometer models used, manufacturers making these models, software analytical tools used, and specific terminology such as those involved in gating.

To cluster the articles, a univariate statistic at the article level was generated for each data type. First, the TF-IDF matrix of terms was created using the abstract, methods, or full text of the set of PLOS articles. These TF-IDF matrices were filtered on a variable number of terms identified for a data type and multiplied by the l2 cosine similarity of each term’s word embedding vector to that data type’s word embedding vector to increase the weight of more similar terms. The resulting weights from each set of terms were summed together for each article to generate an article level “termFreq” statistic. We varied the number of nearest terms as well as data type word embedding vectors (and thereby the terms themselves) by changing the set.seed parameter of the data type generator function. We could then identify the effect of such changes on recall of a positive control set when leveraging the derived “termFreq” statistic for clustering.

1000 random PLOS articles were sampled repeatedly using bootstrap (n = 5) and a set of positive control articles for each data type was added to each sample. These positive control articles were identified using the rplos and rentrez packages. The rplos package was used to search for articles mentioning words within the figure title or caption of results that would indicate generation of a data type whereas the rentrez package was used to identify PLOS articles having an entrez link to the SRA database, which would indicate deposition of sequencing data (see methods section). These positive control articles should have a greater potential to generate the corresponding data type and contain the terminology indicating this; therefore, we would expect our clustering approach to identify a larger proportion of these positive controls. One might also expect that using a larger number of terms for a data type would increase the recall ability of the clustering to detect the signal within the positive control.

The articles were clustered into two clusters: one containing a lower mean “termFreq” statistic and one containing a higher mean “termFreq” statistic for each data type. The cluster with the higher mean “termFreq” was compared to the positive control fraction specific to that data type to calculate the recall (those in the high cluster that were also positive control) (Fig. 3). The results demonstrate that using a greater number of terms tended to result in a higher recall of the positive control articles for most data types. Interestingly, certain data types (e.g. flow cytometry) did not show an improvement or had a decrease in recall, which would suggest that increasing the number of terms leads to a plateau on which no improvement is gained or the signal itself may be diluted by other data types. The plateau might be reached with fewer terms for certain data types like flow cytometry. To test this hypothesis, we obtained the list of top terms for each data type and leveraged their GloVe word embedding to visualize a variable number of top terms in a lower number of dimensions with the T-SNE algorithm (Fig. 4). Interestingly, we observed that using a lower number of terms resulted in each data type having a highly distinct cluster in the lower dimensional embedding. Increasing the number of terms resulted in a gradual overlap in terms that spanned multiple data types supporting that there is an optimal number of terms for each data type that balances precision and recall by including relevant terms and avoiding ambiguous terms.

**Fig. 3.**
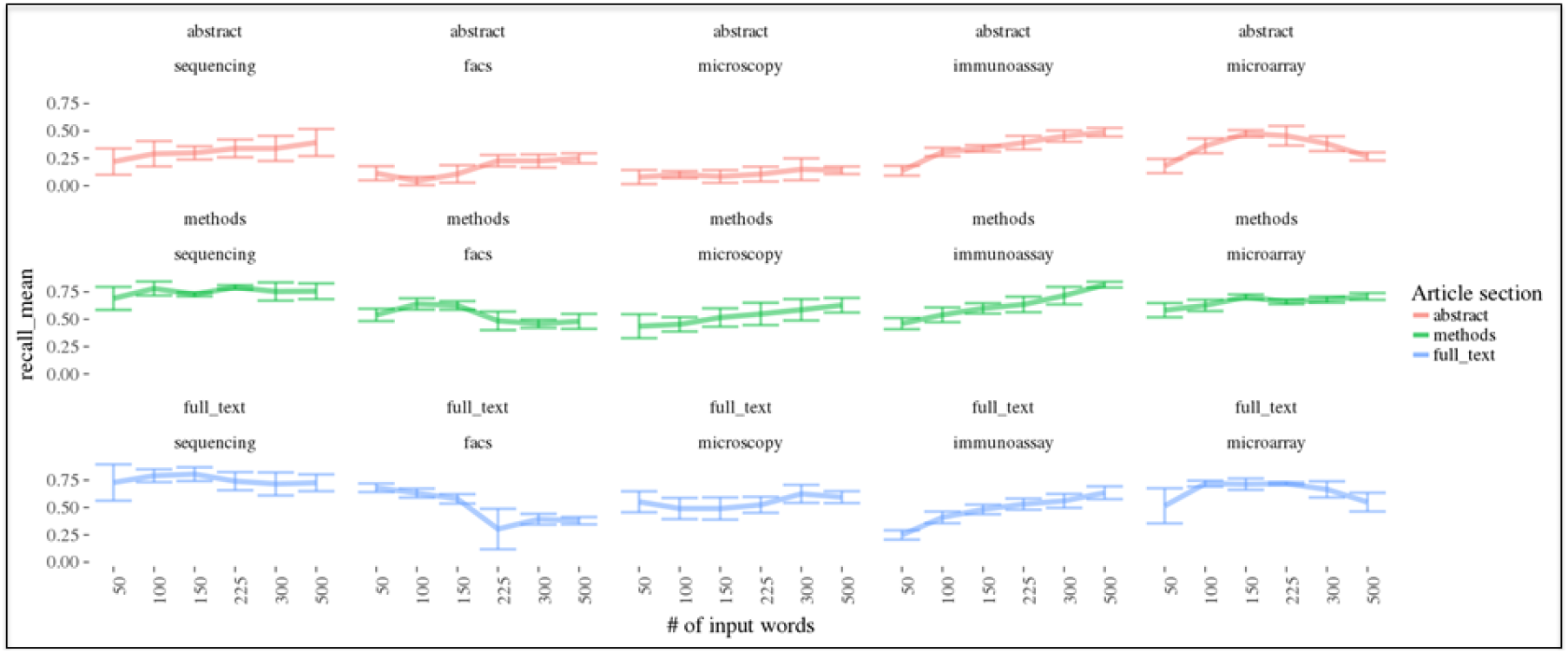
Optimizing number of terms to use on positive control. The effect on recall of varying the number of nearest terms to a data type vector when clustering positive control articles is shown. This representative result is shown for the data type word vector generated by the function having an input random seed start of 3. 5 different sets of 1000 random articles were added to a sample of positive control articles for each data type (sequencing = 93, facs = 200, microscopy = 200, immunoassay = 200, microarray = 73 respectively), the “termFreq” statistic was calculated at the article-level, and the set of “termFreqs” for the articles in each bootstrap sample was clustered using the Ckmeans.1d.dp algorithm. The cluster containing the higher mean “termFreq” was compared to the positive controls to determine the fraction of positive control identified (the recall). The results for each data type and article section used for the clustering when varying the number of input terms is compared illustrating that the use of the methods or full text generally results in a higher recall rate across data types and # of terms used for clustering.

**Fig. 4.**
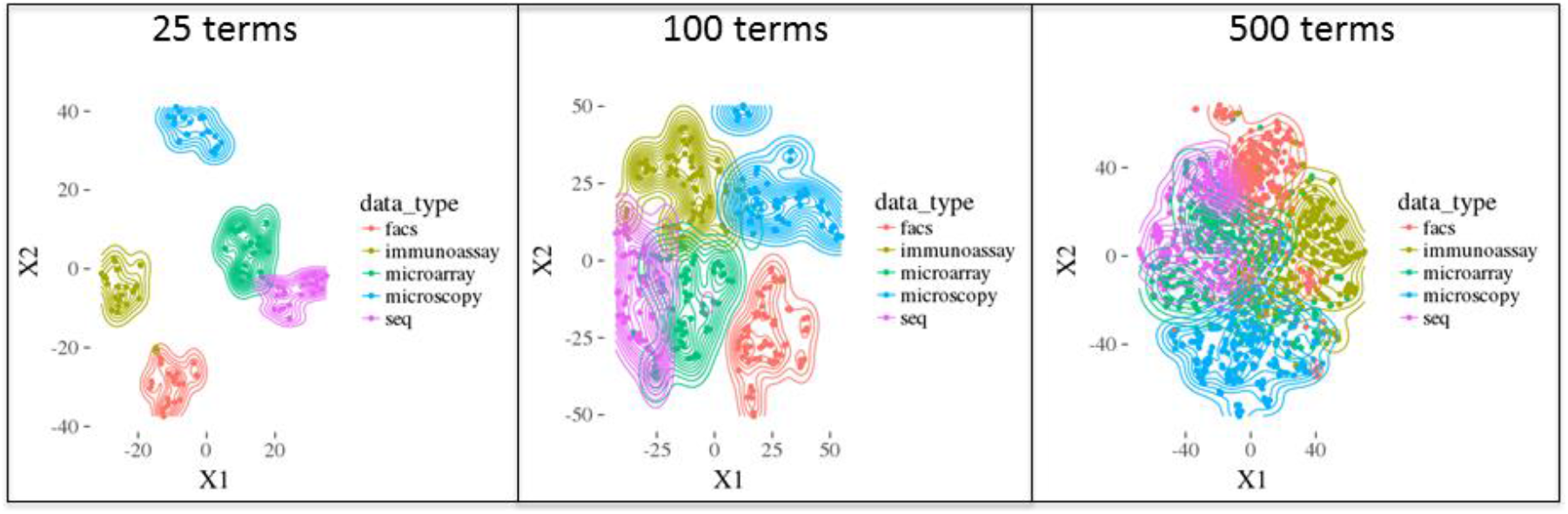
Visualizing increasing term usage similarity via t-SNE. To give an illustration of the relationship of the top terms for each data type, the top terms via l2 normalized cosine similarity to the five biomedical data type word vectors using optimized approach in Table 1 is shown. The word vectors for these terms were visualized using the t-SNE algorithm (using the Barnes-Hut approximation) to embed the high-dimensional vector relationships between the set of words into 2 dimensions (X1 and X2). The resulting visualization captures word similarities such that words often used in the same context within a methods section are nearer to each other. Each color represents the corresponding terms for each of the five biomedical data types analyzed and the three plots correspond to visualization of an increasing number of terms. When the number of terms are increased, there is a greater overlap in the number of words either spanning more than one data type or near words from other data types. This is especially apparent in data types measuring similar outputs such as the expression of genes (e.g. sequencing and microarray).

We also observed that the use of the methods and full text resulted in higher recall rates of the positive control articles. This makes intuitive sense given that most the technical language is in the methods and results sections and not the abstract; therefore, the signal would be more strongly associated with these two sections. The results support that, until the plateau is reached, the added terms identify articles using less common terminology indicative of a data type. A disadvantage to the inclusion of these additional terms might be a decrease in the resulting precision of positively identified articles because there would be a greater potential for ambiguous terms corresponding to multiple data types. These ambiguous terms are observed within similar data types; for instance, there is an overlap between the sequencing and microarray data types (Fig. 4) as these measure gene expression and genomic variation albeit using distinct technologies.

To test the stochastic nature of the function generating each data type vector, we determined the mean recall for the bootstrap samples mentioned above using 5 different set.seed parameters for the function. If stochastic effects do not have a large impact on the resulting data type vectors and terms that are produced, then the “termFreq” statistic would not vary greatly and the clustering would perform similarly regardless of set.seed parameter. The mean recall for a bootstrap sample prepared using different seeds would not show a high standard deviation between samples in this case. Fig. 5 shows a plot of the average of different mean recall values from 5 different set.seed parameters using variable numbers of terms and the specified article section. In most cases, the standard deviation of the mean recall is not large suggesting that the stochastic effects are minimal and result in similar “termFreq” statistics between bootstrap samples. Where there are larger standard deviations, it is when the abstract is used. This may be explained by the lack of technical and methodological terminology used in abstracts as well as the greater variation between abstracts. Some abstracts include language on the key outcomes and results while others omit such language. As stochastic effects produce slight variations in the terms included in the final data type signatures used to produce the “termFreq”, the abstracts may be more likely to be affected.

**Fig. 5.**
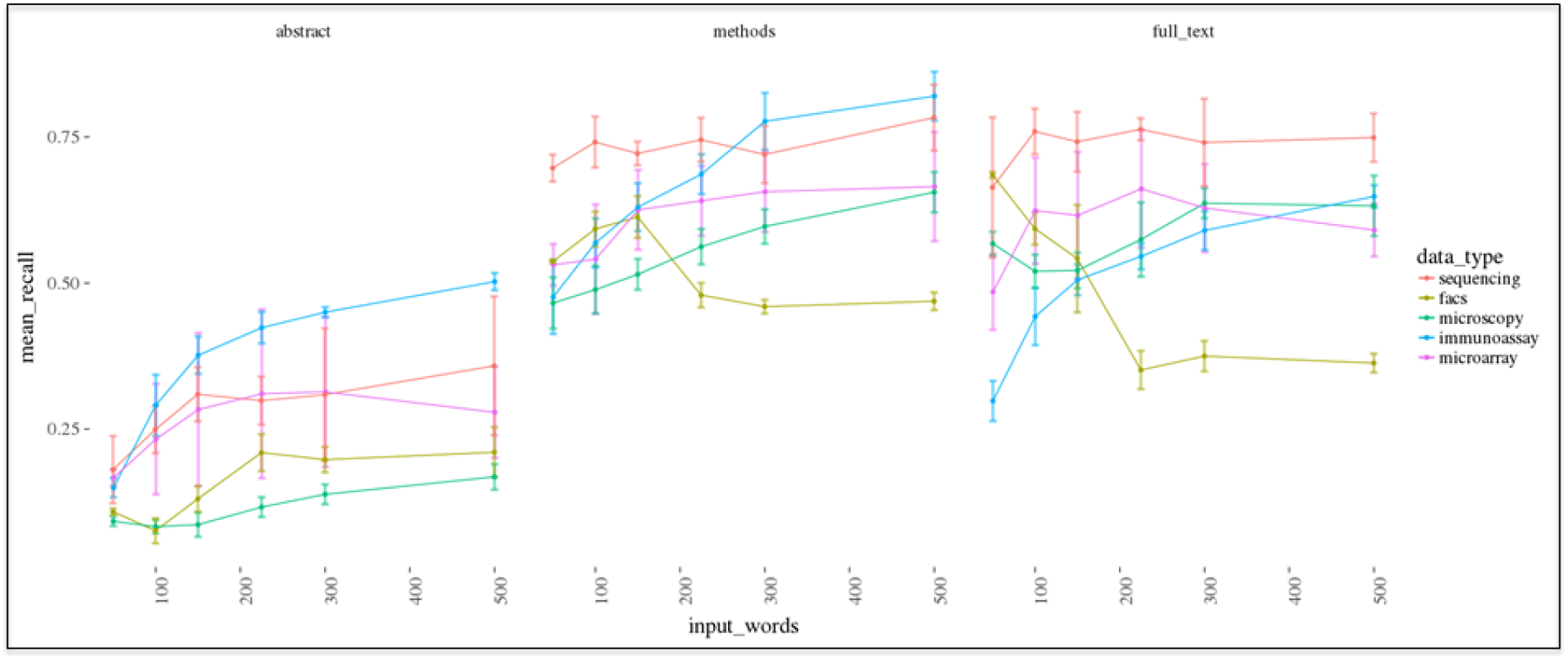
Recall when varying random start of function that generates data type vector. The standard deviation between mean recall when varying the random seed start (1 to 5) from the bootstrap samples on the positive control is shown in the graph above. The trend when varying number of input terms for clustering of the abstract, methods, or full text can be seen. 5 different sets of 1000 random articles were added to a set of positive control articles for each data type (sequencing = 93, facs = 200, microscopy = 200, immunoassay = 200, microarray = 73 respectively), the “termFreq” statistic was calculated at the article-level, and the set of “termFreqs” for the articles in each bootstrap sample was clustered using the Ckmeans.1d.dp algorithm. The cluster with the higher mean “termFreq” was compared to the positive controls to determine the fraction of positive control identified (the recall). The mean recall was calculated for each bootstrap experiment (n = 5) representing the five random seed values tested and the average of these mean values was plotted along with the standard deviation to illustrate how the stochastic effect of changing the random seed affects the resulting recall on the positive control. For most data types and article sections analyzed, the resulting mean values do not show large standard deviations.

### Validation of approach using manually annotated articles

The use of positive control permitted us to test whether signal could be captured in terms of recall but not the quality of such signal with regards to the specificity of the data type of interest. We were not able to determine whether addition of terms might dilute the “termFreq” statistic for a data type and result in a higher chance of imprecision when clustering. To test this, we manually annotated 177 articles likely to be enriched in the five data types we are testing by sampling NIAID funded and positive control articles. The following data types were represented at the following levels via annotation in the manually annotated articles: sequencing (20/177), microarray (60/177), flow cytometry (87/177), immunoassay (62/177), and microscopy (87/177) which was at an equivalent or greater level to the fraction of articles corresponding to each data type within the entire PLOS set of articles we analyzed. We annotated articles based upon whether technologies being used or samples being used could result in the production of a data type. If the methods section or figures mentioned these technologies or samples, then the paper was annotated as producing that data type.

We used bootstrap sampling (n = 5) of 1000 random PLOS articles which were added to the 177 manually annotated articles to test the effect on precision and recall of clustering when varying the number of terms used to generate the “termFreq” statistic. Increasing the number of terms resulted in a decrease in precision with corresponding increase in recall in most cases (Fig. 6). As we observed in the analysis of the positive controls, we observed that adding flow cytometry terms decreased the recall of the manually annotated articles. This provides support to the notion that additional terms are beneficial until a plateau is reached in which more terms results in decreased recall or precision (Fig. 4). Empirical testing is required when generating the “termFreq” statistic as the number of terms to balance precision and recall varied for each data type.

**Fig. 6.**
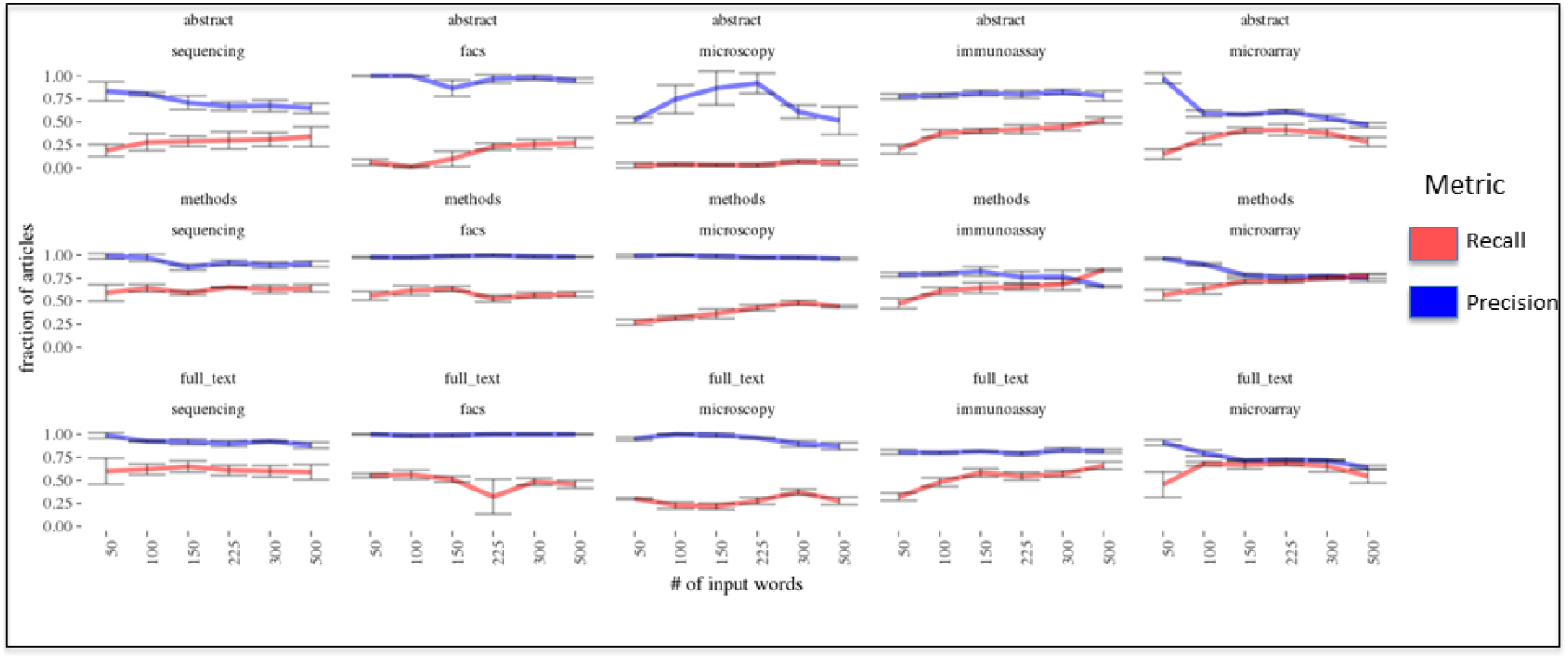
Precision and recall when varying the number of terms. The effect on precision and recall of varying the number of nearest terms to a data type vector when clustering manually annotated articles is shown. A representative result is shown (the other input seeds are similar and can be obtained in the supplementary materials) for the data type word vector generated by the function having an input random seed start of 3. 5 different sets of 1000 random articles were added to a set of manually annotated articles for each data type (n = 177 articles containing the following data types: sequencing = 20 articles, facs = 87 articles, microscopy = 87 articles, immunoassay = 62 articles, microarray = 60 articles), the “termFreq” statistic was calculated at the article-level, and the set of “termFreqs” for the articles in each bootstrap sample was clustered using the Ckmeans.1d.dp algorithm. The cluster containing the higher mean “termFreq” was compared to the manually annotated articles to determine the fraction correctly identified. The results for each data type and article section used for the clustering when varying the number of input terms is compared illustrating that, in general, the use of the methods or full text generally results in a higher recall rate and greater precision across data types.

Using the methods section or full text tends to result in a better balance of precision and recall as compared to the abstract (Fig. 6). The methods section provided the best balance. This makes sense given that the methods and full-text would contain the sections of the article with the technical language indicating whether a data type would be produced. The abstract might have this information depending upon the journal but often omits technical language or simply does not contain enough technical terms to derive a meaningful signal. It was nevertheless exciting to note that the use of the abstract could detect production of each data type in the annotated articles with high precision albeit low recall. Still, the number of terms to use was highly variable and impacted by the stochastic variance in the generator function (supplementary information). While the results are promising, the analysis is a proxy for how the clustering is performing and would benefit from a larger validation set.

To test the stochastic effect of the generator function, we determined the mean recall and mean precision for the bootstrap samples on the annotated articles mentioned above using 5 different set.seed parameters for the generator function. Fig. 7 shows a plot of the average of these different mean recalls from the 5 different set.seed parameters. In most cases, the standard deviation of the mean recall is minimal and is like what we observed for the analysis of the positive controls. This supports that our manual annotation is correctly identifying each data type. It also strongly suggests that the resulting **“termFreq” statistic is not** significantly affected by the stochastic nature of the generator function. For the precision analysis shown in Fig. 8, we observed that the use of abstract resulted in larger standard deviations in mean precision when varying set.seed parameter. Lack of terminology as well as the variation in the use of such technical terminology between abstracts likely explains this higher standard deviation. Most sampling resulted in a good balance of precision and recall (Figs. 7 and 8).

**Fig. 7.**
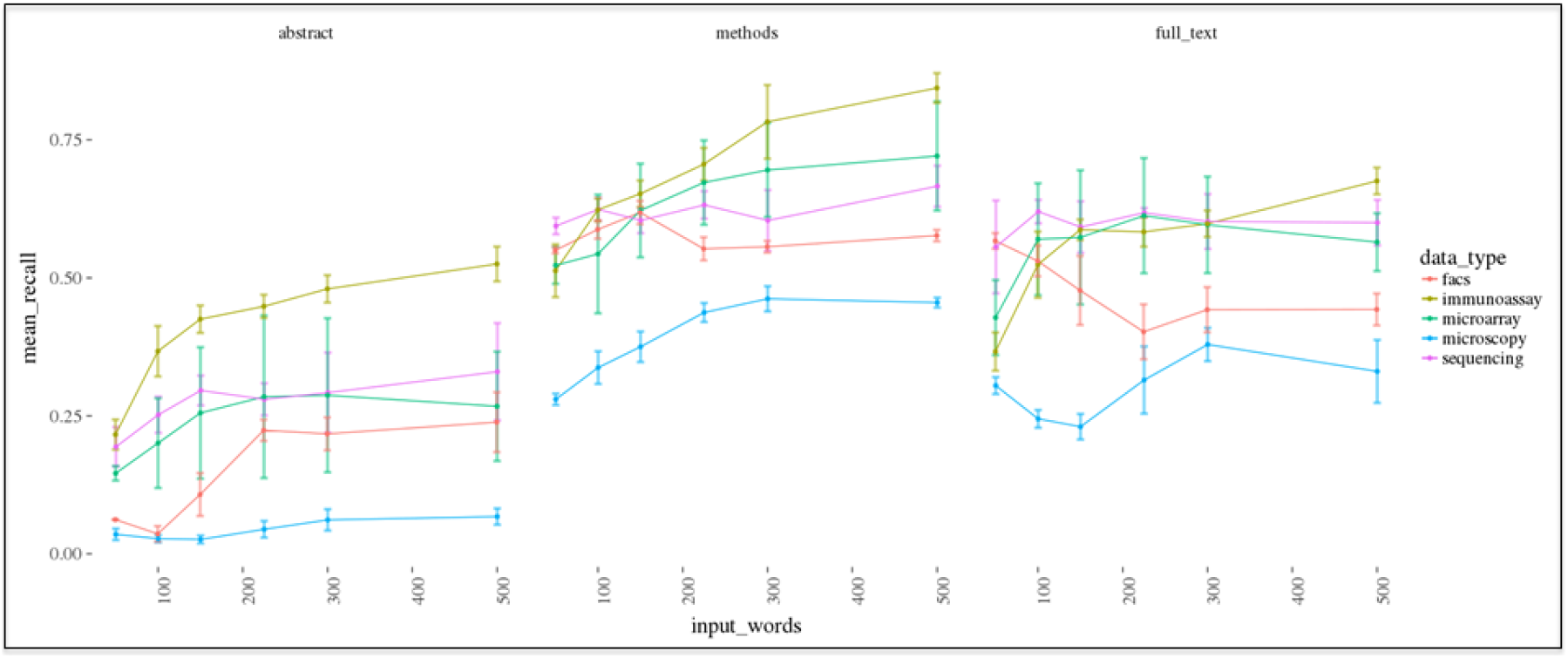
Recall when varying random start of function that generates data type vector. The standard deviation between mean recall when varying the random seed start (1 to 5) from the bootstrap samples on the manually annotated articles is shown in the graph above. The trend when varying number of input terms for clustering of the abstract, methods, or full text can be seen. 5 different sets of 1000 random articles were added to a set of manually annotated articles containing each data type (n = 177), the “termFreq” statistic was calculated at the article-level, and the set of “termFreqs” for the articles in each bootstrap sample was clustered using the Ckmeans.1d.dp algorithm. The cluster with the higher mean “termFreq” was compared to the manually annotated articles to determine the fraction of manually **annotated articles identified (the recall). The mean recall was calculated for each** bootstrap experiment (n = 5) representing the five random seed values tested and the average of these mean values was plotted along with the standard deviation to illustrate how the stochastic effect of changing the random seed affects the resulting recall on the manually annotated articles. For most data types and article sections analyzed, the resulting mean values do not show significant standard deviations.

**Fig. 8.**
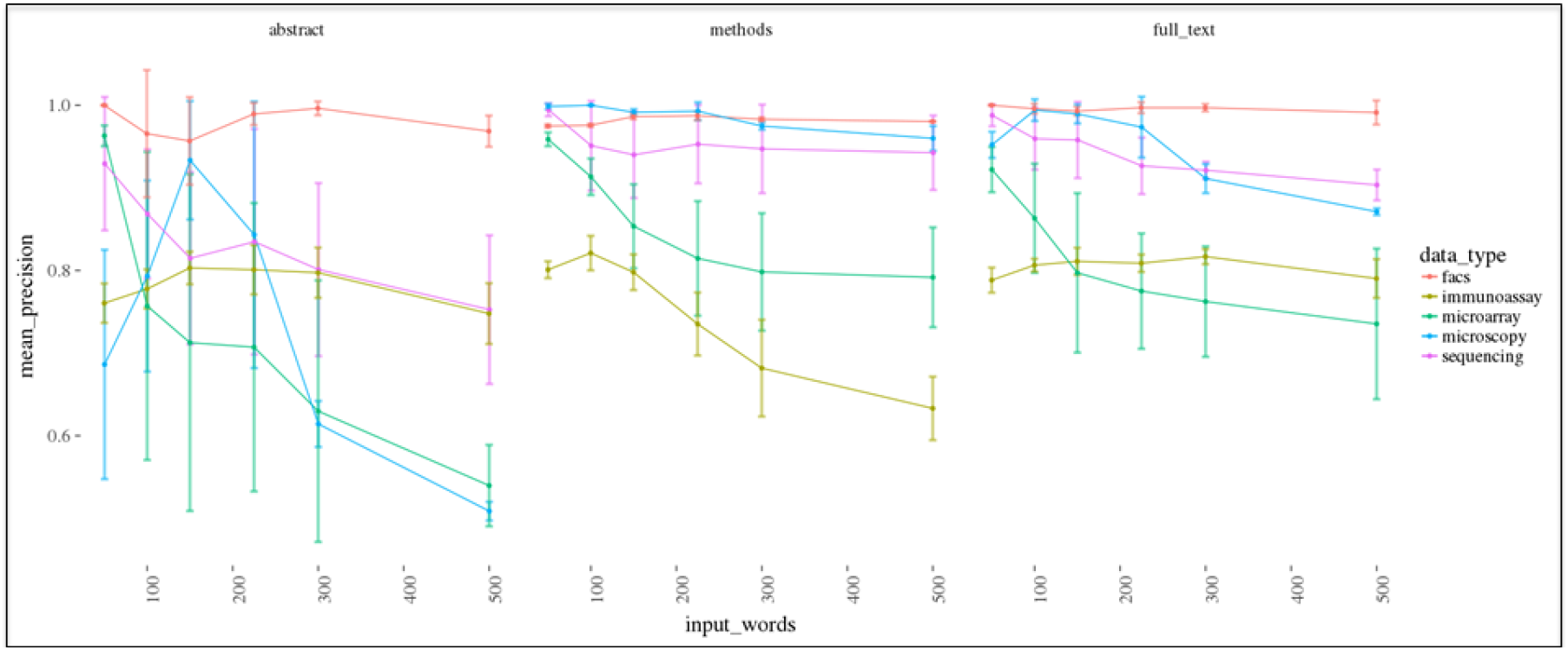
Precision when varying random start of function that generates data type vector. The standard deviation between mean precision when varying the random seed start (1 to 5) from the bootstrap experiments on the manually annotated articles is shown in the graph above when varying number of input terms for clustering of the abstract, methods, or full text. 5 different sets of 1000 random articles were added to a set of manually annotated articles containing each data type (n = 177), the “termFreq” statistic was calculated at the article-level, and the set of “termFreqs” for the articles in each bootstrap sample was clustered using the Ckmeans.1d.dp algorithm. The clusters were compared to the manually annotated articles to determine the fraction of manually annotated articles identified correctly (precisions). The mean precision was calculated for each bootstrap experiment (n = 5) representing the five random seed values tested and the average of these mean values was plotted along with the standard deviation to illustrate how the stochastic effect of changing the random seed affects the resulting precision on the manually annotated articles. For most data types and article sections analyzed, the resulting mean values do not show significant standard deviations although the use of the abstract did show higher standard deviations for microarray and sequencing data types.

### Estimating the data being produced in PLOS articles

We determined the optimal clustering approach by analyzing the manually annotated bootstrapping results to identify a set of parameters that resulted in a useful balance between precision and recall. We used the optimal clustering approach (Table 1) to estimate the data types generated by PLOS articles over time from 2004 onwards (Fig. 9). This analysis lets us understand the dynamics in the biomedical data ecosystem. Interestingly, we observed a sharp decrease in the proportion of microarray data being produced with a stable generation of sequencing data (it dipped around 2008 and has recently begun a gradual increase). Such an observation would be consistent with the technological developments taking place in genomics in the past decade. We noted that microscopy and flow cytometry data generation has fluctuated over time (between 15-25% and 5-9% respectively). The proportion of immunoassay data increased around 2007 along with a similar increase in flow cytometry. Interestingly, this is around the same time that PLOS added “PLOS Pathogens” and “PLOS Neglected Tropical Diseases” to its list of supported journals. These two journals support publications exploring immunological and infectious disease research that would be expected to generate these two types of data. The raw number of data types being produced each year increased until around 2013 and then remained level. Finally, we analyzed PLOS articles indicating NIAID funding (n = 6, 357) and compared them to PLOS articles not indicating NIAID funding (n = 116, 919). We hypothesized that the NIAID funded articles would show a significant increase in representation of immunology data types such as flow cytometry and immunoassay. We observed an increased proportion of NIAID funded articles generating flow cytometry and immunoassay data (~ 3 and 2-fold respectively) when comparing the two groups (Fig. 10).

**Fig. 9.**
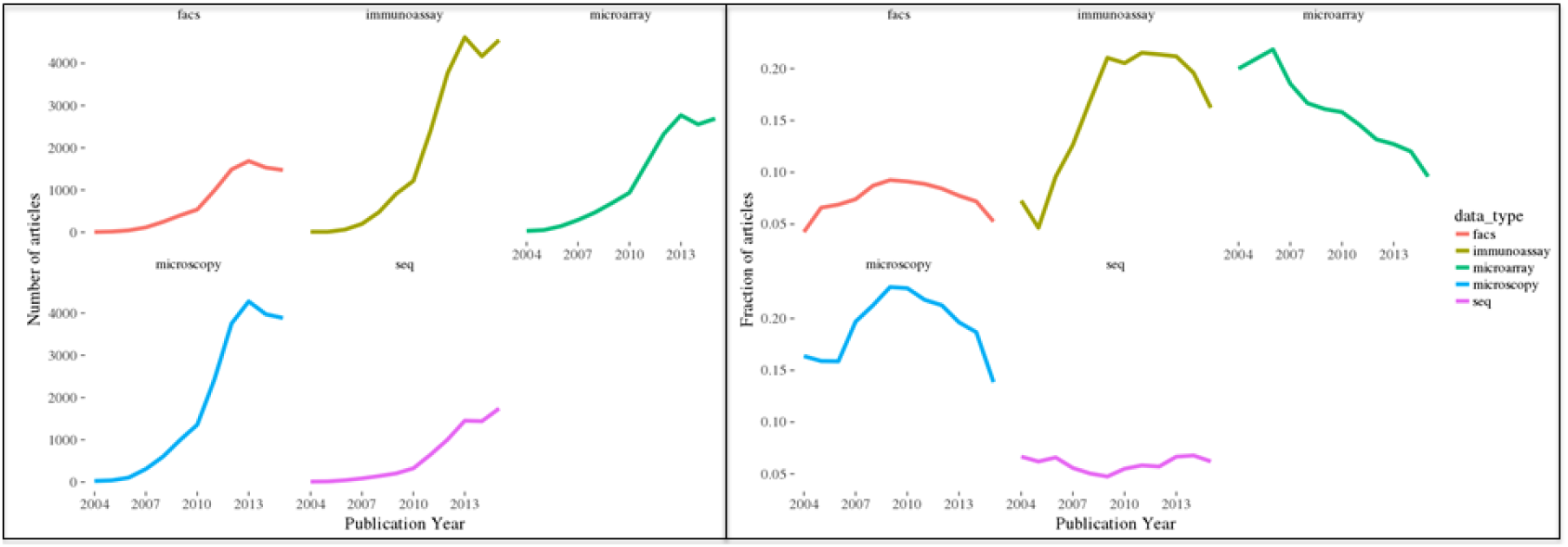
Estimating PLOS articles generating each biomedical data type over time. PLOS articles published after 2004 (n = 129888) were clustered using the optimal clustering approach in Table 1 for each data type. An estimate of the fraction of articles published each year generating each data type as determined using the clustering approach is shown. The left panel shows the raw number of articles producing each data type whereas the right panel shows the proportion of PLOS articles for a given years producing each data type. The raw number of each data type increased steadily until ~2013 and then remained relatively flat. One interesting trend is the gradual decrease in the proportion of microarray data being produced with corresponding increase in sequencing data being produced after 2008. There is also a fluctuation in the fraction of microscopy and flow cytometry data according to the estimate. Interestingly, the proportion of immunoassay data increased from ~2005 onwards eventually plateauing. PLOS Pathogens and PLOS Neglected Tropical Diseases were added to the PLOS journals in 2005 and 2007 respectively, which could correspond with the increase of immunoassay data at that time as well as the similar increase in flow cytometry data (albeit at a lower level).

**Fig. 10.**
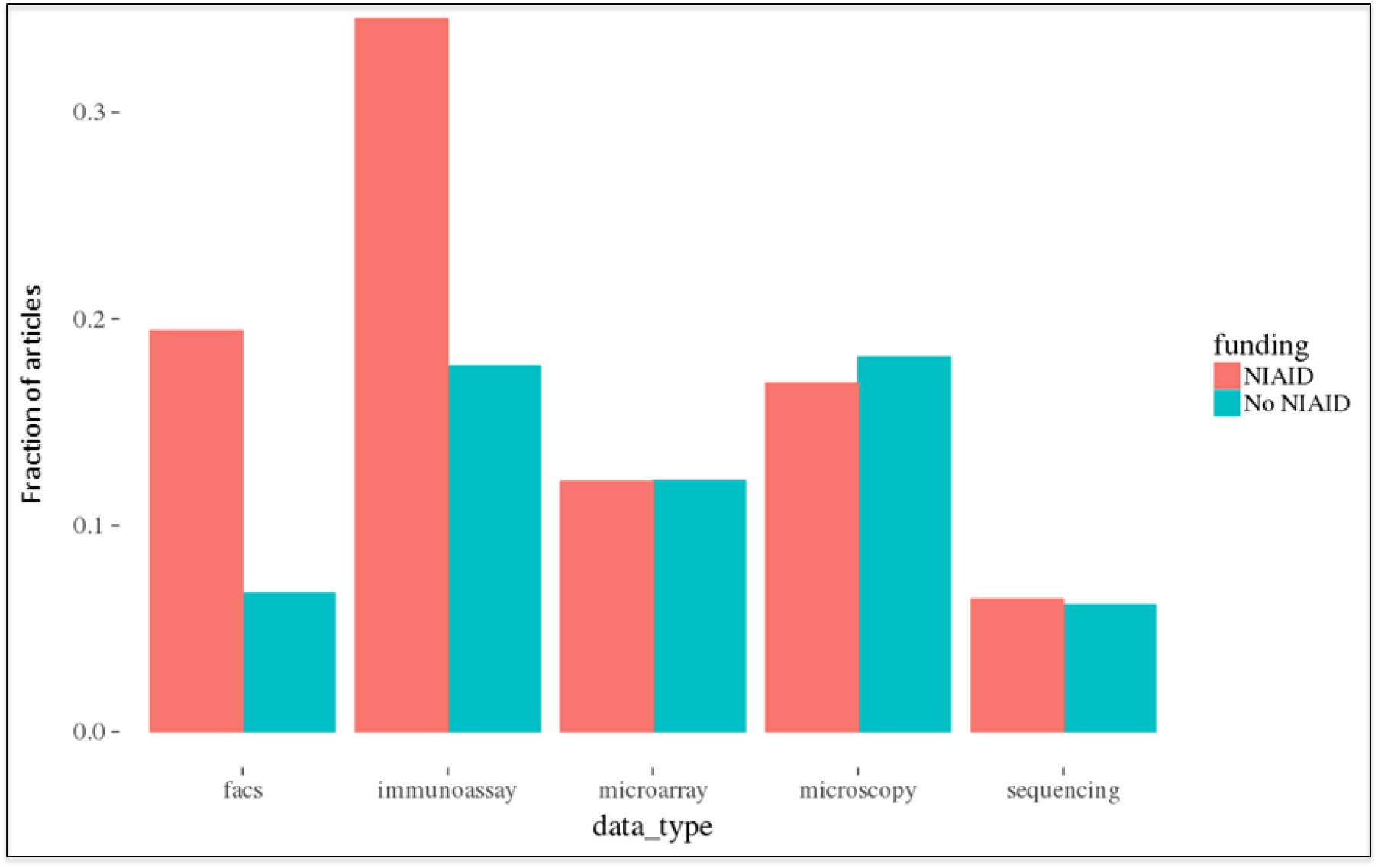
Estimating fraction of NIAID-funded PLOS articles generating each data type. PLOS articles indicating NIAID funding (n = 6325) or articles indicating no NIAID funding (n = 116, 919) were clustered using the optimal clustering approach for each data type in Table 1. An estimate of the fraction of articles published each year generating each data type as determined using the optimal clustering approach is shown. Both flow cytometry and immunoassay data showed a higher fraction of articles in the NIAID-funded subset than in articles not indicating NIAID funding. Other data type fractions were at comparable levels.

**Table 1:** Optimal clustering approach. The table shows the optimal parameters identified for clustering of each data type by balancing the trade-off between mean precision and recall on the manually annotated articles (n = 177) as tested by bootstrap sampling in Figs. 7 and 8. In parenthesis are the standard deviations from five bootstrap samples.

| Data Type | Random seed | Article Section | # of terms | Mean Precision | Mean Recall |
| --- | --- | --- | --- | --- | --- |
| Sequencing | 2 | Methods | 500 | .89 (+/- .027) | .73 (+/- 0.027) |
| Microarray | 4 | Methods | 500 | .76 (+/- .035) | .77 (+/- .02) |
| Flow Cytometry | 3 | Methods | 150 | .989 (+/- .01) | .63 (+/- .027) |
| Immunoassay | 4 | Methods | 225 | .75 (+/- .033) | .72 (+/- .037) |
| Microscopy | 5 | Methods | 300 | .98 (+/- .0009) | .48 (+/- .018) |

## Conclusions

The results demonstrate how to use the GloVe algorithm and the free text of biomedical research articles to identify useful signal for statistical analysis of data production. For the five data types we tested, representing the diversity of the biomedical data being produced, the approach could reliably detect positive control articles as well as provide a useful balance of precision and recall when tested on manually annotated articles. Such an approach could be scaled up to the entirety of articles within PubMed to estimate the amounts and types of data being produced to better quantify this critical research output and to facilitate greater transparency and reproducibility by aiding in the identification of the primary datasets used to reach a scientific conclusion. Another promising use of this approach is to observe other trends or insights of interest (e.g. the emergence of technologies, secondary analysis approaches, or new research areas).

Our estimates from the 129, 918 PLOS articles analyzed indicate that significant proportions of these articles are producing flow cytometry, immunoassay, microscopy, microarray, and sequencing data (~7%, 19%, 18%, 12%, and 6% respectively). These proportions are likely low estimates (especially for microscopy data) given the mean recall observed for the annotated articles (Table 1). Our analysis of 129,918 PLOS articles published between 2003 and 2016 assess that 59,543 articles generated roughly 81, 407 data sets consisting of one of the five biomedical data types we tested. Of those data-producing articles, ~ 1.4 data types were produced per article (supplementary information). Despite the limited scope in terms of data types and publications examined, the analysis demonstrates that a vast amount of biomedical data is being generated. The broader diversity of biomedical articles publishing a greater variety of biomedical data would be expected to generate significantly more.

NIH is the largest biomedical research agency in the world and serves as an interesting example for data production. A search of PubMed to identify NIH funded research articles, excluding reviews, published in 2016 returned 91, 685 articles. Were a comparable scale of biomedical data being produced from this set of articles, then roughly 40,000 of these articles would be expected to generate approximately 56, 000 datasets consisting of one of the five data types we analyzed. For a wider perspective on the production of data, we identified 1, 160, 334 research articles published in PubMed in 2016 by structuring a query to eliminate review articles. If the same proportions of these articles were generating the five data types we analyzed then roughly 81000, 220000, 140000, 210000, and 70000 would produce flow cytometry, immunoassay, genomic microarray, microscopy, and high-throughput sequencing data respectively. These observations highlight a different sort of scientific productivity beyond the traditional bibliographic metrics that are normally quantified. Identification of data being produced from research articles can provide additional context on a work’s impact, give proper credit to an investigator producing the data, enhance transparency and reproducibility, and inform stakeholders in the biomedical data ecosystem about how the ecosystem is developing over time.

## Supporting Information

S1 Supporting.7z

S1 Supporting.7z. This file contains the documentation and source code to generate the figures and results presented in this article. It also contains supplementary figures and supporting information.

## Author Contributions

**Conceptualization:** DL GR

**Methodology:** DL GR

**Data Curation:** GR

**Formal analysis:** GR

**Investigation:** GR

**Project administration:** DL GR

**Software:** GR

**Validation:** GR

**Visualization:** GR

**Writing – Original Draft Preparation:** GR

**Writing – Review & Editing:** DL GR

## Acknowledgements

The findings and conclusions of this report are those of the authors and do not necessarily represent the views of the National Institute of Allergy and Infectious Diseases, National Institutes of Health, or the United States Government. The authors would like to acknowledge helpful feedback and discussion with Quan Chen, Fenglou Mao, Dan Rotrosen, and Charles Hackett.

